# Myofibril orientation as a metric for characterizing heart disease

**DOI:** 10.1101/2021.09.24.461697

**Authors:** Weikang Ma, Henry Gong, Vivek Jani, Maicon Landim-Vieira, Maria Papadaki, Jose R. Pinto, M. Imran Aslam, Anthony Cammarato, Thomas Irving

**Author notes:** Corresponding author: Thomas Irving.

## Abstract

Myocyte disarray is a hallmark of many cardiac disorders. However, the relationship between alterations in the orientation of individual myofibrils and myofilaments to disease progression has been largely underexplored. This oversight has predominantly been due to a paucity of methods for objective and quantitative analysis. Here we introduce a novel, less-biased approach to quantify myofibrillar and myofilament orientation in cardiac muscle under near physiological conditions and demonstrate its superiority as compared to conventional histological assessments. Using small-angle X-ray diffraction, we first investigated changes in myofibrillar orientation at increasing sarcomere lengths in permeabilized, relaxed, wildtype mouse myocardium by assessing the angular spread of the 1,0 equatorial reflection (angle σ). At a sarcomere length (SL) of 1.9 μm, the angle σ was 0.23±0.01 rad, decreased to 0.19±0.01 rad at a SL of 2.1 μm, and further decreased to 0.15±0.01 rad at a SL of 2.3 μm (p<0.0001). Angle σ was significantly larger in R403Q, a MYH7 hypertrophic cardiomyopathy (HCM) model, porcine myocardium (0.24±0.01 rad) compared to WT myocardium (0.14±0.005 rad, p<0.0001) as well as in human heart failure tissue (0.19±0.006 rad) when compared to non-failing samples (0.17±0.007 rad, p=0.01). These data indicate that diseased myocardium suffers from greater myofibrillar disorientation compared to healthy controls. Finally, we showed that conventional, histology-based analysis of disarray can be subject to user bias and/or sampling error and lead to false positives. Our method for directly assessing myofibrillar orientation avoids the artifacts introduced by conventional histological approaches that assess myocyte orientation and only indirectly evaluate myofibrillar orientation, and provides a precise and objective metric for phenotypically characterizing myocardium. The ability to obtain excellent X-ray diffraction patterns from frozen human myocardium provides a new tool for investigating structural anomalies associated with cardiac diseases.

**Statement of Significance:** We introduce a precise and quantitative approach to directly measure myofibrillar and myofilament orientation in cardiac muscle under near physiological conditions as a novel tool for phenotypically characterizing striated muscle systems. We use this technique to demonstrate that myocardium from disease model organisms and failing human myocardium suffers from greater myofibrillar disorientation compared to healthy controls. We also demonstrate that excellent diffraction patterns can be obtained from frozen and thawed human myocardium. Given the ready availability of frozen human heart tissue in tissue banks, this capability opens up a large space of potential experiments relating sarcomere structure to dysfunction in cardiac disorders.

## Introduction

A great deal of effort has gone into elucidating the molecular interactions involved in thick and thin filament-based regulatory mechanisms of force production by the sarcomeres of cardiac muscle, which, when perturbed, can trigger disease (1–6). Much less scrutiny, however, has been given to the very basic question of how myofibrillar and myofilament level alignment may be affected by discrete pathologies and how such changes could impact myocardial function. Classically, myocyte disarray, defined here as the degree to which adjacent myocytes are oriented either obliquely or perpendicularly to each other, is a hallmark of hypertrophic cardiomyopathy (HCM) (7, 8). HCM is further characterized by increased myocardial wall thickness, hyperdynamic contractile properties, impaired energy balance, incomplete myocyte relaxation and diastolic dysfunction (9, 10). While HCM can lead to heart failure (HF), determining HCM-associated HF incidence is problematic due to substantial etiological and clinical heterogeneity (11). Nonetheless, population studies have estimated that more than 50% of patients diagnosed with HCM present HF symptoms (12, 13). In the majority of these individuals, HF manifests with normal systolic function and diastolic dysfunction from left ventricular hypertrophy, with a minority developing HF with reduced ejection fraction (HFrEF).

HFrEF is defined by an ejection fraction of 40% or less, comprises roughly 50% of HF cases, and is characterized by cardiomyocyte loss due to ischemia, mutation, myocarditis, or valvular disease (14). However, electron microscopy (EM) analysis of myocardial tissue in a canine model of heart failure revealed disrupted myofilament structure and loss of regular filament lattice structure likely contributed to depressed force production (17). Unfortunately, microscopy-based methods require painstaking analysis of tissue in a conditional manner, and cannot resolve other critical information such as cross-bridge formation and quantification of structural changes with variable calcium concentrations and is more subject to sample error given the amount of tissue analyzed. These pitfalls, among others, provide an impetus for developing additional approaches which allow for an accurate, unambiguous, and objective analysis of myofilament and myofibrillar disarray in both preclinical models of HF and in human disease, particularly HFrEF.

Small-angle X-ray diffraction of cardiac tissue is a uniquely powerful technique to assess sarcomere structure under near physiological conditions. The thick filaments in the sarcomeres of vertebrate muscle are packed into a hexagonal lattice with interdigitated thin filaments in the trigonal positions. The equatorial X-ray diffraction pattern (Figure 1) arises from the projected density of the mass in the A-band of the sarcomere. The 1,0 equatorial reflection arises from thick filaments, while the 1,1 equatorial reflection arises from both thick and thin filaments. The ratio of the intensity of the 1,1 equatorial reflection to that of the 1,0 equatorial reflection (I_1,1_/I_1,0_) is a measure of the relative degree of association of myosin heads with thin filaments, both under resting conditions (1, 18–20) and during contraction (19–23). The distance between the 1,0 equatorial reflection to the beam canter reflections can be used to calculate the interfilament lattice spacing, d_10_. There is more information in the equatorial pattern, however, in the form of the angular spread of the equatorial reflections relative to the equatorial axis. In striated muscle, the equatorial reflections are perpendicular to the myofibrillar longitudinal axis; thus, the angular spread of the equatorial reflections across the equator (Figure 1) will be a direct measure of the degree of alignment of the sarcomeres relative to the long axis of the preparation. The angular spread of X-ray reflections has previously been used to characterize the relative orientation of collagen fibers in connective tissue (24, 25). However, this approach has not previously been used, to our knowledge, to assess the alignment of contractile machinery in cardiac tissue.

**Figure 1.**
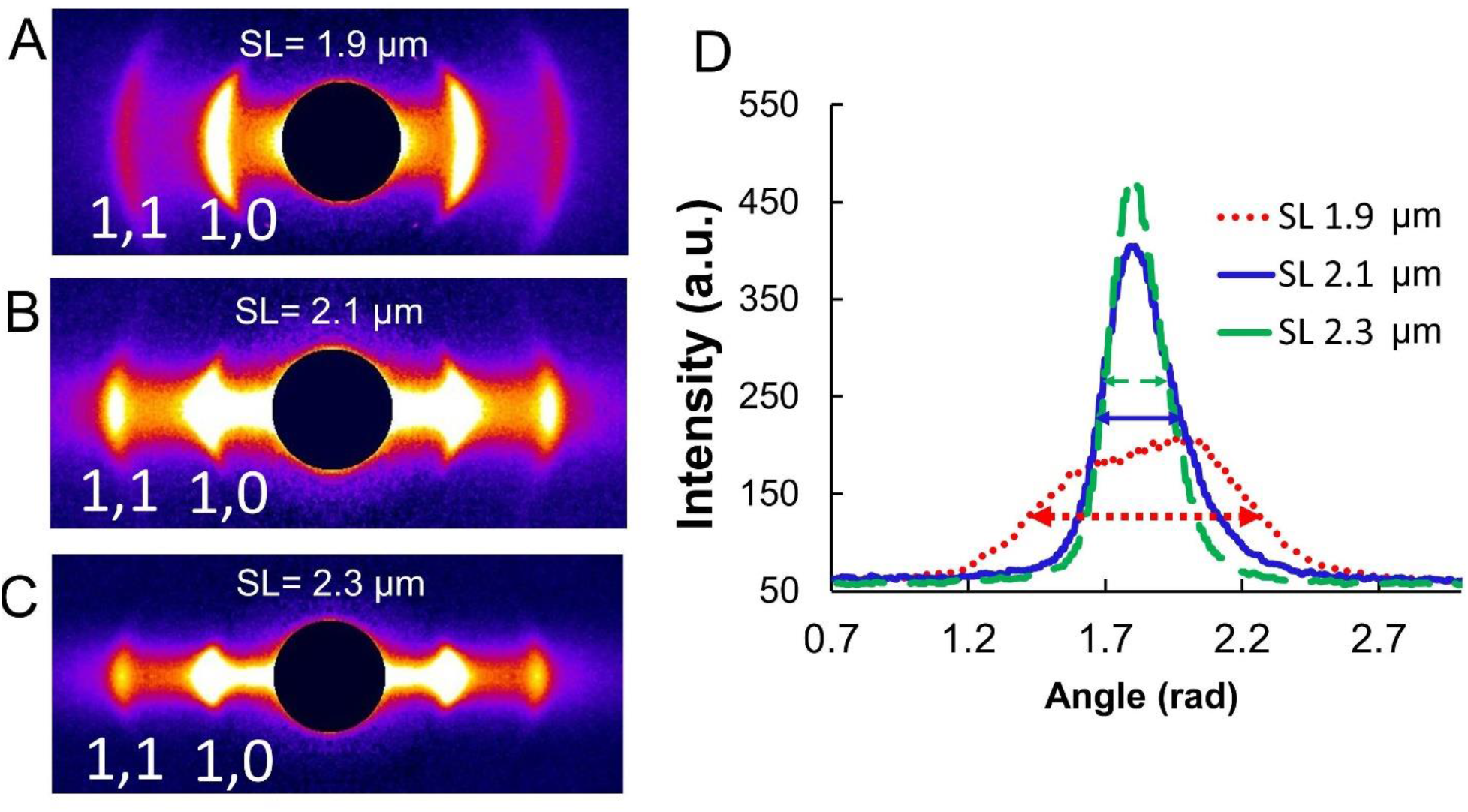
Equatorial X-ray diffraction patterns from permeabilized murine myocardium. Representative equatorial X-ray diffraction patterns from permeabilized mouse myocardium (A, B & C) and the angular intensity profile of the 1,0 equatorial reflection (D) at different sarcomere lengths. The full width at half maximum (~2.36 σ) of the peaks is indicated by the double headed arrows at corresponding colors.

Here, we first investigated the changes in myofibril structure with increased sarcomere length in mouse myocardium with the motivation of evaluating a fundamental sarcomere geometric mechanism that could help explain myofilament length-dependent activation (LDA), the phenomenon where increasing sarcomere length results in additional force for the same calcium concentration. Indeed, we observed improved myofibril alignment with increased sarcomere length in normal mouse myocardium, consistent with LDA. This observation prompted us to attempt to validate our technique’s ability to quantify myofibril-level disarray and correlate this to cardiac disease. As a first step towards this goal, we studied myofibrillar disorientation in transgenic porcine myocardium expressing R403Q mutant myosin. The R403Q mutation was the first identified HCM-causing mutation in β-cardiac myosin (26), and hearts from R403Q transgenic mice showed increased myocyte disarray (27). By measuring the angular spread of the equatorial reflections, we quantitatively confirmed that, in large animal models, R403Q myocardium also suffers from a higher degree of myofibrillar disorientation as compared to control. We then extended our studies to human tissue, demonstrating, for the first time, that high quality 2D X-ray diffraction patterns, suitable for detailed structural analysis, can be obtained from frozen human heart tissue. Myofibrillar disorientation was quantified in right ventricular myocardium from non-failing donors (Non-Failing) vs. patients with HFrEF that was not preceded by HCM. We show that the HFrEF tissue displayed significant myofilament- and myofibril-level disorientation relative to Non-Failing controls.

Our method for assessing myofibrillar orientation provides an objective measurement of alignment by sampling all myofibrils exposed to the X-ray beam and avoids sampling errors and artifacts due to tissue processing associated with conventional histological approaches. The ability to provide precise quantitation of myofibrillar orientation offers a new metric for phenotypically characterizing these tissues and may provide insight into depressed force production observed in human heart failure. X-ray diffraction has been shown to provide insights into the relationship of disease related mutations to sarcomere-level structural changes in animal model systems (6, 28–31) and our new findings indicate that this technique can now also be applied to human myocardium. Together, these findings establish X-ray diffraction as a uniquely powerful tool for establishing the relation of myofibrillar structure to function in healthy and diseased myocardium across animal models and human patients.

## Methods

### Mouse myocardium preparations

All procedures using live mice were done in accordance with protocols approved by the Institutional Animal Care and Use Committees (IACUCs) of the Florida State University and the Illinois Institute of Technology. Papillary muscles were dissected from mice and permeabilized with 1% Triton X in relaxing solution (containing in mM: 6.3 Na_2_ATP, 6.48 MgCl_2_, 10 EGTA, 100 BES, 10 phosphocreatine, 49.76 KPropionate, 10 DTT, and creatine kinase 10 U/ml) overnight. The muscles were washed with fresh cold relaxing solution and muscles were further dissected into fiber strips, clipped on aluminum T-clips and stored in cold relaxing solution for the day’s experiments.

### Porcine myocardium preparation

The data from the porcine myocardium reported here were from a re-analysis of the patterns reported by Anderson et al. (32).

### Human myocardium preparation

Human subjects were enrolled under protocols approved by the Institutional Review Boards at the Johns Hopkins University, the University of Pennsylvania, and the Gift-of-Life Donor Program of Pennsylvania (33). Human myocardial tissue was procured as described previously (33). Briefly, failing hearts (n = 5) were obtained via explantation at the time of orthotopic heart transplantation, while non-failing hearts (n = 5) were obtained from brain-dead organ donors. Non-failing hearts were not used for transplantation due to donor age. Hearts were rapidly arrested using high-potassium cold-cardioplegia and open-chest surgical excision, then transported in cold Krebs-Henseleit Buffer (KHB) on wet ice to an on-campus laboratory. Tissue was then rapidly dissected, snap-frozen in liquid nitrogen, and stored at −80°C until transport. RV septal tissue from patients showing low RV Fmax (15, 16) was used in all RV studies.

Frozen human RV tissue were permeabilized as previous described for porcine myocardium (32). Briefly, pieces of frozen human RV tissues (3mm^3^) were put in relaxing solution (containing in mM: 100 BES, 10 EGTA, 6.6 MgCl_2_, 10 creatine phosphate, 15 creatine kinase, 6.2 ATP, 5 NaN_3_, pH 7.0) containing 1% Triton X-100 at room temperature for 2-3 hours. The skinning solution was changed every hour. The muscle was then washed with relaxing solution prior to application of aluminum T-clips and stored in cold relaxing solution for the day’s experiments.

### Histology preparation and measurement of myocyte alignment

Conventionally prepared hematoxylin and eosin (H/E) and Masson’s Trichrome stained tissue were obtained from both non-failing and failing human right ventricular septal myocardium. Myofibrillar disarray was quantified using an image processing-based approach, described in detail elsewhere (27, 34). Briefly, between 4-6 cropped 1000 × 1000-pixel sections of longitudinally sectioned RV myocardium at 20x magnification were identified for analysis from both H/E and Masson’s trichrome stained sections for each patient. A longitudinally sectioned region was considered suitable for analysis if there was minimal fibrosis, few histologic artifacts, and minimal spaces/cracks within the tissue. Identified regions were exported as “.png” images and imported into MATLAB (version R2018b, Natick, Massachusetts: The MathWorks Inc, 2020). A 2D fast Fourier transform was applied to each image, and a power spectral density was determined as described (34). Next, for each angle, a histogram of the distribution of the relative number of myocytes aligned at each angle *θ* was determined, and the percent myocyte alignment, quantified as the relative proportion of myocytes within 20° of the mean myocyte alignment angle, was determined for each image.

### Small-Angle X-Diffraction

Equatorial X-ray diffraction patterns were collected from freshly permeabilized muscle strips using the small-angle instrument on the BioCAT beamline 18ID at the Advanced Photon Source, Argonne National Laboratory (35). The X-ray beam was focused to ~0.06 × 0.15 mm at the detector plane. The sample-to-detector distance was ~3.5 m and the X-ray wavelength was 0.103 nm. Isolated cardiomyocyte bundles (~200 µm diameter, 2-3 mm long) were mounted between a force transducer (Model 402A, Aurora Scientific) and a static hook. Force was monitored using Muscle Dynamic Control system (Model 610A, Aurora Scientific). Sarcomere length was adjusted by laser diffraction using a 4-mW HeNe laser. Diffraction patterns were collected at sarcomere lengths of 1.9 μm, 2.1 μm and 2.3 μm for mouse myocardium experiments and 2.0 μm for human myocardium experiments. X-ray exposures were 1 s at an incident flux of ~3×10^12^ photons per second, and the patterns were collected on a CCD-based X-ray detector (Mar 165; Rayonix Inc. Evanston, IL).

### X-ray Data Analysis

The data were analyzed using data reduction programs from the MuscleX software package developed at BioCAT (36). The angular divergence of the 1,0 equatorial X-ray reflections was measured by the “Scanning Diffraction” routine in the MuscleX software package. Briefly, the routine obtains 2D and 1D radially integrated intensities of the pattern. The 1D radially integrated intensity trace was fit assuming Gaussian profiles as a function of radial spacing for the diffraction peaks to calculate the standard deviation (width σ) of the peak intensity distribution in the radial direction. In this process, the routine obtains the integrated intensity of each equatorial reflection as a function of the integration angle. Gaussian profiles are fit to the projected peak intensities to find the standard deviation of the orientation angle (angle σ) to calculate the angular divergence.

### Statistics

Statistical analyses were performed using GraphPad Prism 9 (Graphpad Software). The results are given as mean ± SEM unless otherwise stated. One-way paired ANOVA with Geisser-Greenhouse correction and Tukey multiple comparison test were used to compare columns in figure 2 in a pairwise manner. Two-tailed unpaired Mann-Whitney tests were used for the data from porcine and human myocardium shown in figure 3 and figure 4. Symbols on figures: ns: *p*>=0.05, *: *p*<0.05, **: *p*<0.01, ***: *p*<0.001 and ****: *p*<0.0001.

**Figure 2.**
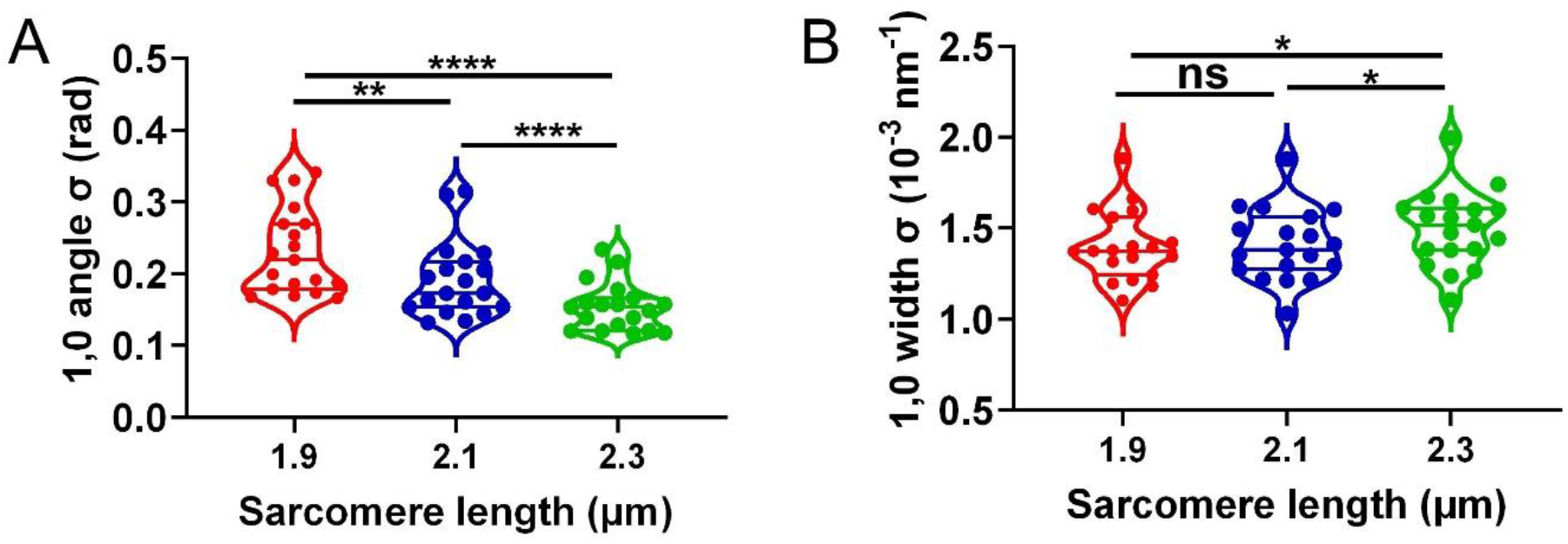
1,0 equatorial reflections in permeabilized mouse myocardium at different sarcomere lengths. **A.** The angular standard deviation of 1,0 equatorial reflections (angle σ) from permeabilized mouse myocardium as a function of sarcomere length. **B**. The standard deviation of the 1,0 equatorial reflections in radial direction (width σ) from permeabilized mouse myocardium as a function of sarcomere length (ns: *p*>=0.05, *: *p*<0.05, **: *p*<0.01 and ****: *p*<0.0001).

**Figure 3.**
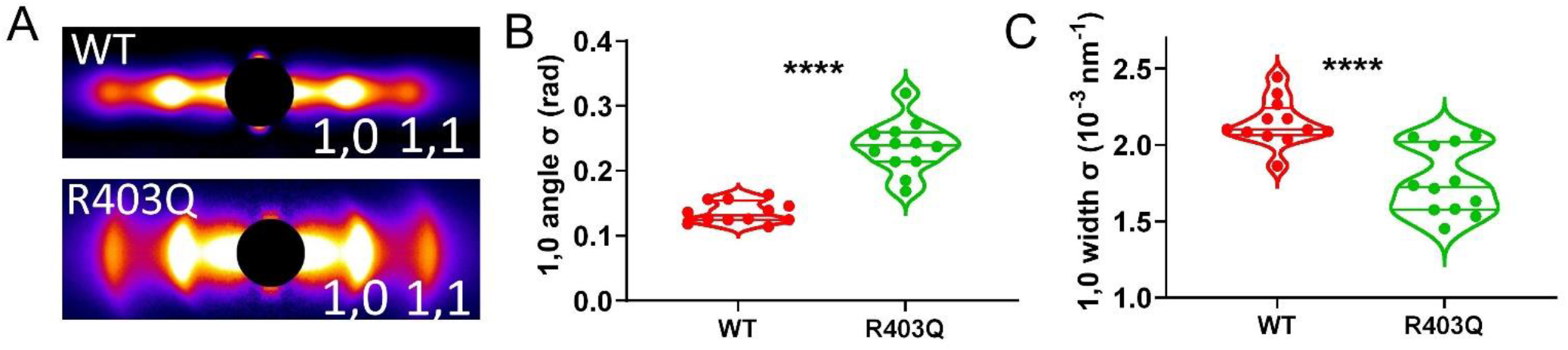
Myofibrillar orientation from permeabilized porcine myocardium. **A,** Representative equatorial X-ray diffraction patterns from permeabilized WT and R403Q porcine myocardium. **B,** The angular standard deviation of 1,0 equatorial reflections (angle σ) from permeabilized WT and R403Q porcine myocardium. **C**, The standard deviation of the 1,0 equatorial reflections in radial direction (width σ) from permeabilized WT and R403Q porcine myocardium (****: *p*<0.0001).

**Figure 4.**
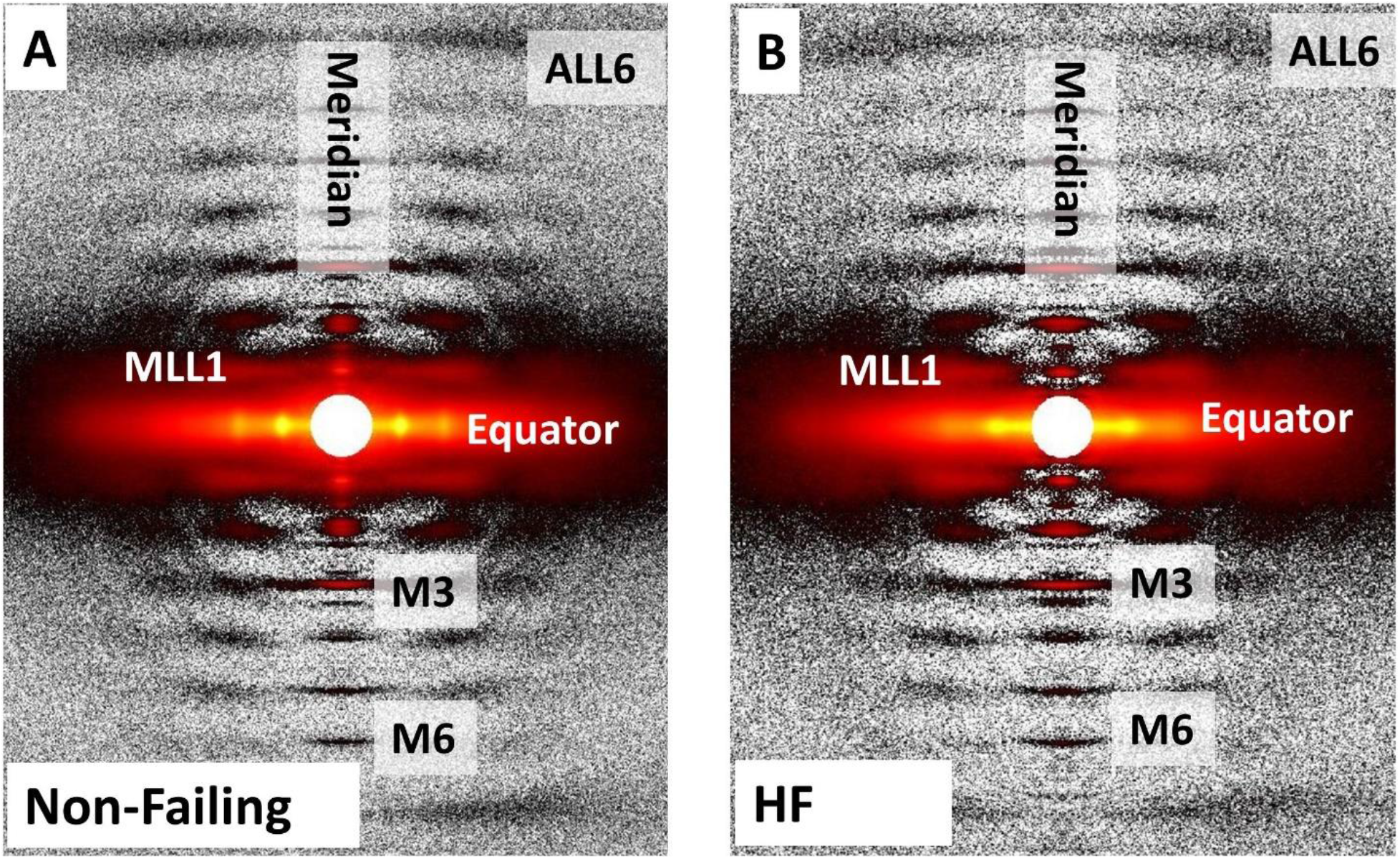
Representative X-ray patterns from frozen non-failing (Non-Failing, panel A) and heart failure patients (HF, panel B) human myocardium. The 1,0 and 1,1 equatorial reflections, the third (M3) and sixth (M6) order myosin-based meridional reflections, first order myosin-based layer lines (MLL1), and the sixth order actin-based layer lines (ALL6) are as labeled.

## Results

To establish a new approach for quantifying the degree of myofilament order, and how it changes with sarcomere length, in native myocardium, we first collected X-ray diffraction patterns from permeabilized WT mouse left ventricle papillary muscle at sarcomere lengths of 1.9 μm, 2.1 μm, and 2.3 μm under relaxing conditions. The equatorial 1,0 and 1,1 reflections were present at all three lengths. The intensity profiles, however, became better defined with increasing muscle sarcomere length, as shown in representative X-ray patterns (Fig 1A, B & C). The equatorial 1,0 and 1,1 reflections were visibly arced at a sarcomere length of 1.9 μm. The azimuthal angular width of the reflections became smaller as the sarcomere length increased, as shown by the angular integrated intensity profiles of the 1,0 reflections in Figure 1D. The standard deviation of the angular projection of the peak (angle σ) was plotted as a function of sarcomere length (Figure 2A), which showed that angle σ decreased by about 20% for every 10% increase in sarcomere length. At a sarcomere length of 1.9 μm, the angle σ for the 1,0 reflection was 0.23 ± 0.01 rad, which decreased to 0.19 ± 0.01 rad at a sarcomere length of 2.1 μm (*p* = 0.005), and further decreased to 0.15 ± 0.01 rad at a sarcomere length of 2.3 μm. (*p* < 0.0001).

The standard deviation of the radial width of the equatorial peaks (width σ) was also calculated as a function of sarcomere length. Width σ for the 1,0 reflection, a measure of heterogeneity in lattice spacing between myofibrils, remained unchanged when the sarcomere length was increased from 1.9 μm (1.40 ± 0.04 (10^−3^ nm^−1^)) to 2.1 μm (1.41 ± 0.04 (10^−3^ nm^−1^); *p* = 0.78). However, width σ significantly increased at a sarcomere length of 2.3 μm (1.50 ± 0.05 (10^−3^ nm^−1^)) when compared to sarcomere lengths of 1.9 μm (*p* =0.04) and 2.1 μm (*p* = 0.04). These data demonstrate that the decrease in angle σ at longer sarcomere length is accompanied by a small but significant increase in lattice spacing heterogeneity in skinned, wild type myocardium.

Our results from normal mouse myocardium raised the question whether these assays of myofilament ordering could be used to characterize myocardium from disease models. Therefore, we analyzed myocardium from a large animal model expressing the well-characterized, HCM-causing R403Q myosin mutation. Myofibrillar orientation was assessed in permeabilized WT and R403Q porcine myocardium at a sarcomere length of 2.0 μm (Figure 3). Qualitatively, the equatorial reflections are arced in R403Q myocardium, while the equatorial reflections are well defined spots in WT myocardium (Figure 3A). The angle σ was significantly higher in R403Q myocardium (0.24 ± 0.01 rad) relative to WT myocardium (0.14 ± 0.005 rad) (p <0.0001) (Figure 3B). In addition, width σ for the 1,0 reflection is significantly larger in in WT myocardium (2.14 ± 0.04 (10^−3^ nm^−1^)) than in R403Q myocardium (1.76 ± 0.06 (10^−3^ nm^−1^)) (p <0.0001) (Figure 3C). These data are consistent with that from previous studies using small rodent models of R403Q myosin-dependent HCM, which likewise showed disarray at the myocyte level (27).

We next examined the possibility of applying our methods to phenotypically characterize myofibrillar and myofilament alignment in normal and diseased human myocardium. If X-ray diffraction patterns could be obtained from cryo-frozen human heart tissue, they could offer a uniquely powerful and transformative tool for examining structure/function relationships in clinically relevant specimens. Figure 4 shows that the structural integrity of the sarcomeres in these preparations was well preserved. In addition to the equatorial reflections addressed in this study, meridional reflections arising from the axial repeats of myofilaments are strong and extend beyond the sixth order myosin-based meridional reflection (M6). Layer lines arising from the quasi-helically ordered array of myosin heads around the thick filament backbone in relaxed muscle are clearly visible. The first myosin layer line (MLL1) is particularly strong.

To investigate myofilament disarray in human HFrEF, we studied the angular and radial profiles of 1,0 equatorial reflection from frozen healthy human myocardium and myocardium from HFrEF patients under relaxing conditions at a sarcomere length of 2.0 μm. Specifically, we chose human HFrEF samples with depressed maximum calcium-activated force from RV permeabilized myocytes as described previously (15). The angle σ was significantly higher in HFrEF myocardium (0.19 ± 0.006 rad) than non-failing (Non-Failing) myocardium (0.17 ± 0.007 rad) (*p* = 0.01) (Figure 5B), indicating a higher degree of myofibrillar disorientation in HFrEF myocardium. The width σ for the 1,0 reflection was significantly higher in HFrEF myocardium (1.35 ± 0.03 (10^−3^ nm^−1^)) than in non-failing myocardium (1.25 ± 0.03 (10^−3^ nm^−1^)) (*p* = 0.04) (Figure 5C), demonstrating a greater degree in dispersion in lattice spacing between myofibrils in HFrEF myocardium.

**Figure 5.**
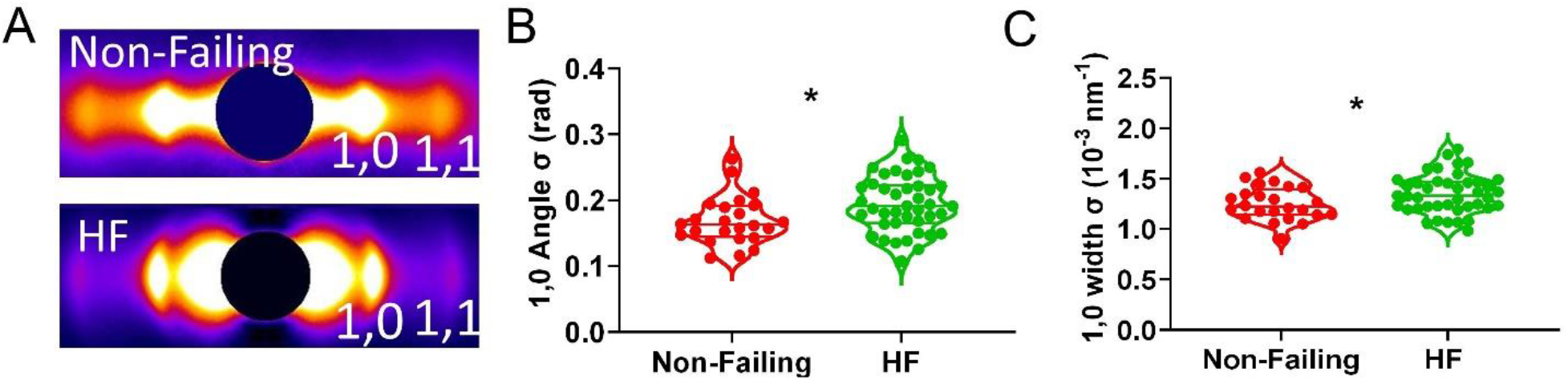
Human myocardium myofibrillar orientation by X-ray from non-failing donors (Non-Failing) and heart failure patients (HF). **A**, Representative equatorial X-ray diffraction patterns from Non-Failing myocardium and HF myocardium. **B**, The angular standard deviation of 1,0 equatorial reflections (angle σ) from permeabilized Non-Failing and HF human myocardium. **C**, The standard deviation of the 1,0 equatorial reflections in radial direction (width σ) from permeabilized Non-Failing and HF human myocardium (*: *p*<0.05,).

Finally, we benchmarked our method for assessing myofibrillar orientation using X-ray diffraction against existing approaches that only assess myocyte orientation, which will be, at best, indirectly related to myobrillar/myofilament alignment. Representative Masson’s trichrome sections from the same non-failing and HFrEF RV samples used for X-ray analysis are shown in Figures 6A and 6B, respectively. Quantification (see Methods) of the percentage of aligned myocytes (Figure 6D) revealed that the percentage of aligned myocytes was significantly reduced in RV HFrEF histologic sections compared to non-failing RV histologic sections. Importantly, these results are consistent with our results showing myofibrillar disarray by X-ray diffraction (Figure 5). Next, to demonstrate the sensitivity of histologic methods to user-based selection criteria, we calculated myocyte alignment in regions *not considered suitable* for analysis in non-failing histologic sections, denoted Non-Failing alt (Figure 6C). This analysis revealed that the percentage of aligned myocytes in these regions was similar to those from HFrEF histological sections (Figure 6D), suggesting that user criteria for histology analysis may be subject to bias and susceptible to false positives.

**Figure 6.**
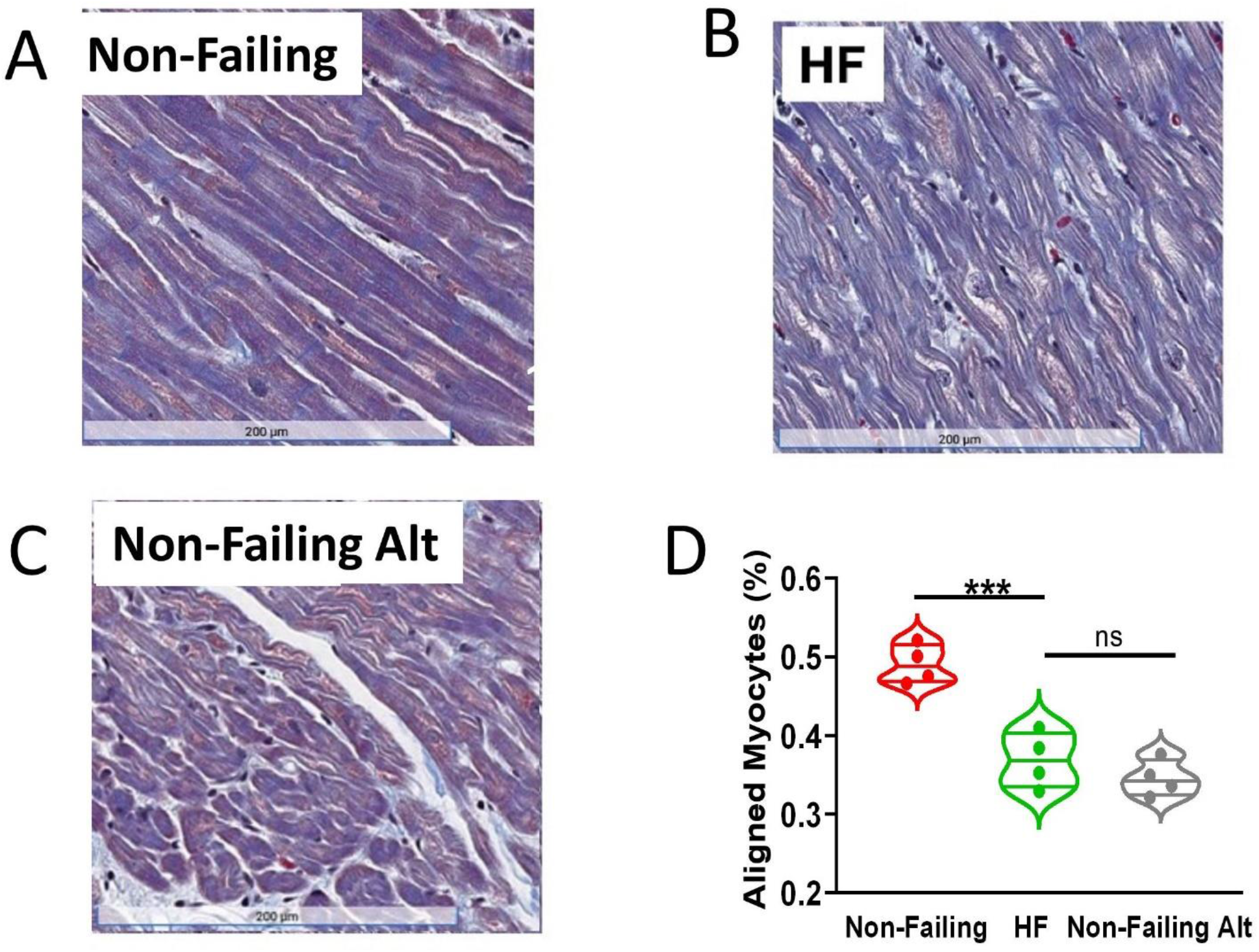
Human cardiomyocyte alignment by histology analysis from non-failing donors (Non-Failing) and heart failure patients (HFrEF). Representative histology images from Non-Failing myocardium (**A**) and HF myocardium (**B**). **C,** Representative histology images from Non-Failing myocardium at an alternative region. **D**, The percentage of aligned myocytes in Non-Failing myocardium, HFrEF myocardium, and Non-Failing myocardium at alternative regions (Non-Failing Alt) that were *not* considered longitudinally aligned or suitable for analysis from histology analysis (ns: *p*>=0.05, ***: *p*<0.001).

## Discussion

### Characterization of myofibrillar orientation in permeabilized murine and porcine myocardium

Myocyte alignment or disarray is commonly used to characterize healthy vs. diseased heart tissue; however, the orientation of myofibrils and myofilaments has been overlooked, largely because the lack of available methods to assess it. Here we quantitatively showed, using X-ray diffraction, that the myofibrils in mouse cardiac muscle cells are relatively poorly aligned at a sarcomere length of 1.9 μm with an angular divergence (angle σ) of 0.24 ± 0.02 rad, and that angular divergence significantly decreased to 0.15 ± 0.01 rad at a sarcomere length of 2.3 μm (Fig 2A). At longer sarcomere lengths, the decrease in angle σ indicates that the myofibrils in cardiac muscle cells are better aligned with the longitudinal axis. When myofibrils contract and produce force, the active force can be separated into both a radial component and an axial component relative to the long axis of the muscle preparation. Better aligned myofibrils in the longitudinal axis could lead to more axial force at longer sarcomere length by redistributing radial components of force to the axial direction and provide a partial explanation for the increased force in LDA.

The radial width σ of the 1,0 equatorial reflection did not change when the myocardium was stretched from 1.9 μm to 2.1 μm. A significant increase in width σ, however, was observed when the sarcomere length was increased to 2.3 μm. The radial width σ is a direct measurement of inter-filament spacing heterogeneity. This inter-filament spacing heterogeneity could come from the spread of lattice spacings among individual myofibrils (inter-myofibrillar) and/or within individual myofibrils (intra-myofibrillar) in the myocytes. An increase in intra-myofibrillar heterogeneity at longer sarcomere length is most likely due to the increase in titin-based passive tension with increasing sarcomere length, being relatively modest between 2.0 and 2.1 μm and increasing exponentially at 2.3 μm and longer sarcomere lengths (37). Titin-based passive tension will have both radial and longitudinal components, that may vary between the myofilaments, but overall tend to compress the lattice and shorten the sarcomere (38). An increase in inter-myofibrillar heterogeneity at longer sarcomere length is likely due to the non-homogeneous transmission of passive force by cytoskeletal components within myocytes such as the desmin intermediate filament network surrounding the myofibrils and inter-connecting the Z-lines (39–41) and the costomeres connecting the Z-lines to the sarcolemma (42, 43). Interestingly, the expression levels of desmin in the heart have been shown to increase in HCM/diastolic dysfunction mouse models (44).

Remodeling of myocyte morphology is a complex process and is regulated by mechanical, hormonal, and hemodynamic stimuli (45). Pathological remodeling of cardiomyocytes is widely observed in end-stage heart failure (46, 47). One of the end results of pathological heart remodeling is cardiomyocyte disarray, which is characteristic of cardiomyopathies including hypertrophic cardiomyopathy (HCM). Myocyte disarray has been proposed to be one of the criteria for the diagnosis of HCM (48, 49). Here, we extend this concept to the myofibrillar level by examining the degree of myofibrillar disorientation in transgenic animal models that are known to display myocyte disarray. Specifically, our results showed that in a porcine model of HCM, with the R403Q mutation, the myocardium displayed a > 70% increase in the 1,0 angular σ compared to WT myocardium. Furthermore, radial width σ of the 1,0 reflection was significantly smaller in R403Q myocardium compared to WT myocardium (Fig 3C), indicating a higher degree of inter-filament heterogeneity. Figure 2B shows that radial width σ and angular σ of the 1,0 reflections are not necessarily coupled, and as such, they can be considered two independent parameters to characterize a given myocardial system.

### Study of frozen human myocardium using X-ray diffraction

Here, we used the methodology introduced in our studies of murine and porcine myocardium to examine the degree of myofibrillar disorientation from failing vs. non-failing human myocardium. Specifically, we investigated frozen explanted heart tissue from a subpopulation of HFrEF patients with known RV myocyte dysfunction, as assessed by depressed maximum calcium-activated tension. Our results demonstrated that myocardium from HFrEF patients displayed larger values of 1,0 angular σ compared to that from non-failing donors and suggests that myofibrillar disarray may contribute to both RV myocyte dysfunction and global RV dysfunction.

The mechanism of depressed sarcomere function in HFrEF as measured by maximal calcium activated tension has been explored in previous studies (15, 16). For example, myofilament disarray has shown to be responsible for diminished force production in a canine model of heart failure (17). Currently, however, the exact mechanisms by which myofilament disarray results in depressed force output are not known, and may not be the only mechanism acting in a given situation. Now that we have a tool to quantify myofilament orientation, experiments can be designed to determine the extent to which myofilament level disarray plays a role in the etiology of any myocardial disorder.

We also show, for the first time, that frozen human myocardium produces high-quality two-dimensional X-ray patterns that may provide valuable information concerning not only the molecular structural basis of the functional behavior in healthy hearts, but also can potentially resolve structural aberrations/correlates that may contribute to depressed or enhanced contractility commonly observed in cardiomyopathies. It has been demonstrated that myosin heads in resting muscle are distributed between the super relaxed state (SRX), characterized by a very low intrinsic ATPase rate, and the disordered relaxed state (DRX) where the intrinsic ATPase rates are much faster (50). The myosin heads in the DRX state are able to participate in cross-bridge formation during contraction while myosin heads in the SRX state are unable to interact with actin but can be recruited by inotropic effectors, including sarcomere stretch, effectively serving as a reserve (51). The equilibrium between myosin heads in SRX and DRX state are important in balancing cardiac physiological functions, such as LDA(51). It has been proposed that many cardiomyopathies are caused by disruption of SRX and DRX equilibria and transitions between these states (52). X-ray diffraction is very sensitive to global structural transitions of myosin heads between SRX and DRX in the context of thick filament based regulatory mechanisms as shown in studies of in rodent and porcine myocardium (3, 32, 53). The ability to obtain this kind of information, in addition to myofibrillar orientation, from human myocardium greatly enhances the value this approach for translational studies as demonstrated in a preliminary report (54). A full report demonstrating increased myofibrillar disarray in myocardium from HFrEF patients will appear elsewhere.

### Comparison to histological measurements

Here, we introduce small-angle X-ray diffraction as a new quantitative method to measure the degree of myofibrillar disorientation in myocardium. X-ray diffraction-based methods have several important advantages over other imaging techniques. X-ray diffraction information may be obtained from permeabilized tissue under near-physiological conditions at a known sarcomere length. The observed X-ray diffraction patterns are the superposition of the individual diffraction patterns from all myofibrils exposed to the X-ray beam, with the azimuthal angular spread of the pattern providing an objective measure of the degree of myofibrillar orientation. Additionally, X-ray diffraction patterns can be taken at the same time as mechanical measurements providing simultaneous structural and physiological information in real-time. Histopathological-based approaches have the advantage of using widely available microscopic tools and well-understood sample preparation protocols. These protocols, however, require multiple steps, including fixation, embedding and microtomy, staining, and image processing, each of which can introduce artifacts (55). None of these preparatory steps are necessary for X-ray diffraction. Furthermore, histological assessments only directly measure myocyte level disarray, and can only indirectly provide inferences into myofibrillar and myofilament level disarray. Here we have shown that myofibrillar orientation is sensitive to sarcomere lengths, and sample shrinkage during any of the histologic processing steps may affect the accuracy of any measurement. Furthermore, with histologic methods, only information encompassed within small regions of the sample selected for imaging and then for analysis are assessed, potentially leading to sampling bias. Histologic methods are also exquisitely sensitive to the choice of the region identified. Regions must contain longitudinally oriented fibers, be free of histologic artifacts, and void of fibrosis and spaces within the tissue. While these criteria can be addressed in animal studies by simply examining more samples, satisfying all criteria may be difficult with histologic sections of where sample material is limiting. Finally, if inappropriate criteria are used for image selection, image processing-based methods may result in a false-positive finding of myocyte disarray, as demonstrated in Figure 6. Given these considerations, histological assessment of myocyte array requires a skilled operator well-versed in fiber architecture.

Nonetheless, the principal advantage of histologic methods is that they can be done anywhere with suitable microscopy facilities, whereas small-angle X-ray diffraction requires the use of a small angle diffraction instrument on a synchrotron X-ray beamline. There are only a small number of such instruments worldwide. In the USA, only the BioCAT beamline 18ID at the Advanced Photon Source routinely does muscle diffraction, but the measurements described here would be feasible on other synchrotron X-ray beamlines worldwide. The ability to obtain small-angle X-ray diffraction patterns from frozen, biopsy-sized, pieces of human myocardium, vastly expands the potential application of this method to assess the contributions of structure and physiology to human heart failure. This provides a motivation for expanding access to synchrotron facilities for studies of human cardiomyopathies, HFrEF, and HFpEF.

Our proposed metric for characterizing myocardium provides otherwise inaccessible information at the myofibril/myofilament level in addition to that from the arsenal of techniques available for phenotypic analysis of diseased myocardium. We anticipate that this will be primarily useful for structurally charactering disease phenotypes and to generate hypotheses that can be tested experimentally to provide biophysical insights into disease etiology. Since this information is available in virtually all X-ray diffraction patterns from muscle, this can be done routinely as part of the analysis of every X-ray diffraction experiment where one is also acquiring lattice spacing and equatorial intensity ratio. Myofibrillar orientation is seen as a complementary tool rather than a replacement for more accessible histological and molecular approaches as well as functional assays that are routinely for characterization cardiomyopathies.

## Conclusions

Small angle X-ray diffraction of permeabilized myocardium can be used as a novel approach to precisely quantify myofibrillar orientation and the distribution of lattice spacings in permeabilized cardiac tissue under near physiological conditions. These metrics can be used to precisely and objectively phenotypically characterize both human biopsies from diseased hearts and experimental systems, including transgenic animal disease models as well as provide biophysical insights into disease etiology. The ability to obtain full two-dimensional X-ray patterns from frozen human myocardium opens up new translational opportunities to relate sarcomere structure to function in health and disease.

## Author Contributions

WM, VJ, HG, MP, and ML designed and performed research. WM, VJ analyzed the data and WM, VJ, JP, IA, AC and TI wrote the manuscript

## Acknowledgments

This research was supported by NIH grants T32 GM73009 (V.J), T32 HL007227 (M.I.A.), HL128683 (J.R.P) and the American Heart Association 19IPLOI34770173 (T.I) and 111POST7210031 (M.P). This research used resources of the Advanced Photon Source, a U.S. Department of Energy (DOE) Office of Science User Facility operated for the DOE Office of Science by Argonne National Laboratory under Contract No. DE-AC02-06CH11357. This project was supported by grant P41 GM103622 and P30 GM138395 from the National Institute of General Medical Sciences of the National Institutes of Health. The content is solely the responsibility of the authors and does not necessarily reflect the official views of the National Institute of General Medical Sciences or the National Institutes of Health.

